# Self-supervised learning for DNA sequences with circular dilated convolutional networks

**DOI:** 10.1101/2023.01.30.526193

**Authors:** Lei Cheng, Tong Yu, Tero Aittokallio, Jukka Corander, Ruslan Khalitov, Zhirong Yang

## Abstract

Due to their intrinsic properties, DNA molecules commonly exhibit long-range interactions along a linear sequence representation. Taking this information into account when modeling DNA sequences is therefore important for obtaining more accurate sequence-based inference. Many deep learning methods have recently been developed for this purpose, but they still suffer from two major issues. First, the existing methods can only handle short DNA fragments, thereby losing longer-range interactions. Second, the current methods require massive supervised labeling while missing most order information within the sequences. Consequently, there is a need to develop an efficient deep neural network modeling framework to extract wide contextual information for more accurate sequence-based inference tasks. Our new framework, named Revolution, takes full DNA sequences as input, without any condensation, and can give accurate predictions for DNA sequences up to 10kbp. In variant effect prediction, our method increases the Area Under the Receiver Operating Characteristics (AUROC) by 19.61% on 49 human tissues on average. Revolution is also demonstrated to work on the plant sequences by improving 2.36% AUROC on average for predicting open chromatin regions (OCRs). The data, models, and code can be freely accessed at https://github.com/wiedersehne/Revolution-DNAPretraining.

## Introduction

With the decreasing cost of DNA sequencing, sequence-based inference has rapidly become popular in many applications, such as taxonomic classification (Rizzo et al., 2015; Yu et al., 2022; Khalitov et al., 2022), enhancer prediction (Sethi et al., 2020; Yang et al., 2017), variant effect prediction (Lee et al., 2015; Avsec et al., 2021), and gene expression prediction (Zhou and Troyanskaya, 2015; Avsec et al., 2021; Kelley et al., 2018; Kelley, 2020). Since the great success of deep learning methods in Natural language processing (NLP), researchers have started to develop computational tools based on deep neural networks for genome sequences, as they are similar to natural language in many aspects (Ji et al., 2021; An et al., 2022).

There are three main research streams for using deep learning techniques for sequence-based inference. Convolutional neural networks (CNNs) are the most popular methods, used in many sequence-based inference tasks, such as gene expression prediction (Avsec et al., 2021; Kelley et al., 2018), regulatory activity prediction (Zhao et al., 2021), and metagenomics gene prediction (Al-Ajlan and El Allali, 2019). For example, DeepMind (Alipanahi et al., 2015) trained a CNN to identify the binding preference of DNA-binding and RNA-binding proteins, and it outperformed previous state-of-the-art computational methods. DeepSEA (Zhou and Troyanskaya, 2015) also applied a CNN to predict the noncoding-variant effects from DNA sequences. Basenji (Kelley et al., 2018) developed a CNN framework to predict cell-type–specific epigenetic and transcriptional profiles in large mammalian genomes from DNA sequences. The second group of methods is recurrent neural networks (RNNs), such as Long Short-Term Memory (LSTM; Hochreiter and Schmidhuber, 1997) and Gated Recurrent Units (GRU; Bahdanau et al., 2014). These methods are also used to capture the sequential characteristic of DNA sequences. For example, Liu et al. (2019) used LSTM to detect DNA base modifications, while Yang et al. (2017) applied GRU for enhancer prediction. Recently, there has been increasing interest in incorporating Transformer or Transformer-like models for sequence-based inference. For example, Khalitov et al. (2022) studied the performances of different X-formers on taxonomy classifications. Wang et al. (2021) presented a transferable Transformer-based method, BindTransNet, for cross-cell type DNA-protein binding prediction. More popularly, researchers tend to combine several deep learning methods to catch the long interactions within the DNA sequences. For example, Enformer (Avsec et al., 2021) first used a CNN tower to condense the sequence to a short piece and then stacked Transformer encoders to integrate long-range interactions. In addition, there are methods that combine CNN and LSTM or GRU for enhancer-promoter interaction (EPI) prediction (Zhuang et al., 2019; Min et al., 2021), DNA binding sites prediction (Zhang et al., 2020), and DNA classification (Gunasekaran et al., 2021). Works on pretraining DNA sequences are also promoted following BERT’s success in NLP (Devlin et al., 2018). DNABERT was a BERT model with a genomic setting (Ji et al., 2021). GeneBERT was proposed for multi-modal pretraining (Mo et al., 2021). MoDNA introduced common DNA functional motifs and domain language in the pretraining (An et al., 2022).

Despite many efforts in applying deep learning to sequence-based inference, the current methods still suffer from one or both of the following issues. First, they cannot scale to very long sequences, which makes it hard to capture long interactions taking place within DNA sequences. For example, the cis-regulatory elements (CREs), cooperating to make use of alternative promoters for biological functions, can be widely spaced. Thus, learning the long interactions within DNA sequences is needed in order to achieve a high-fidelity representation of the underlying biology. Second, the methods without self-supervised learning require much more supervised labels to achieve accurate results. However, it is usually too expensive and time-consuming to acquire sufficiently labeled data for genomic analysis. For example, to identify one enhancer, biologists need to place a fragment of presumed regulatory DNA near a promoter and evaluate whether or not the CRE element increases transcription (Cho, 2012). This task will become super expensive, with nearly one million enhancers present in the mammalian genome. Consequently, there is a need for a model that can generalize well when labeled data is limited.

In this paper, we propose a deep learning architecture called ciRcular dilatEd conVOLUTIONal (Revolution) and apply it to self-supervised learning for long DNA sequences. The Revolution framework includes pretraining and fine-tuning. A circular dilated design of Revolution allows it to capture the long-range interactions in DNA sequences, while the pretraining benefits Revolution with only a few supervised labels. We demonstrate that Revolution can handle long sequences and accurately conduct DNA-sequence-based inference. Experimental results show that Revolution can scale up to 10kbp for pretraining on human genome reference 38 (GRCh38) and plant genome data set. Revolution achieves 91.12% AUROC on a variant effect prediction task in human tissues. On the plant data set, Revolution improves the open chromatin region (OCR) prediction AUROC by 2.36% on average.

## Materials and Methods

### Network Architecture

The key component in our network architecture is convolution with exponentially increasing dilations and circular padding. The dilated convolutions can achieve large receptive fields in a few layers (Oord et al., 2016; Kalchbrenner et al., 2016; Lea et al., 2017; Bai et al., 2018). As shown in Figure 1A, the dilation at the *l*-th layer is d_*l*_ = 2^*l−*1^, and every output element can receive information from all input elements in *O*(log_2_ *N*) layers for a length-*N* sequence. The circular padding convolves signals at one end with the other end (In Figure 1A, the red nodes present signals from the other end). Such a padding scheme relieves the boundary effect and provides robust capacity for data shifting (Kayhan and Gemert, 2020; Alsallakh et al., 2020). Let 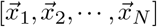 denotes an input sequence of the *l*-th layer. The output sequence 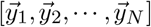 of circular dilated covolution is given by

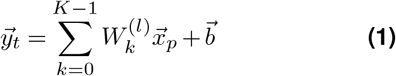

where *t* = 1,…, *N*, 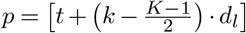 mod *N*, *K* is the kernel size, 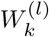 is the *k*-th convolution weights, and 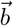 is the bias.

**Figure 1.**
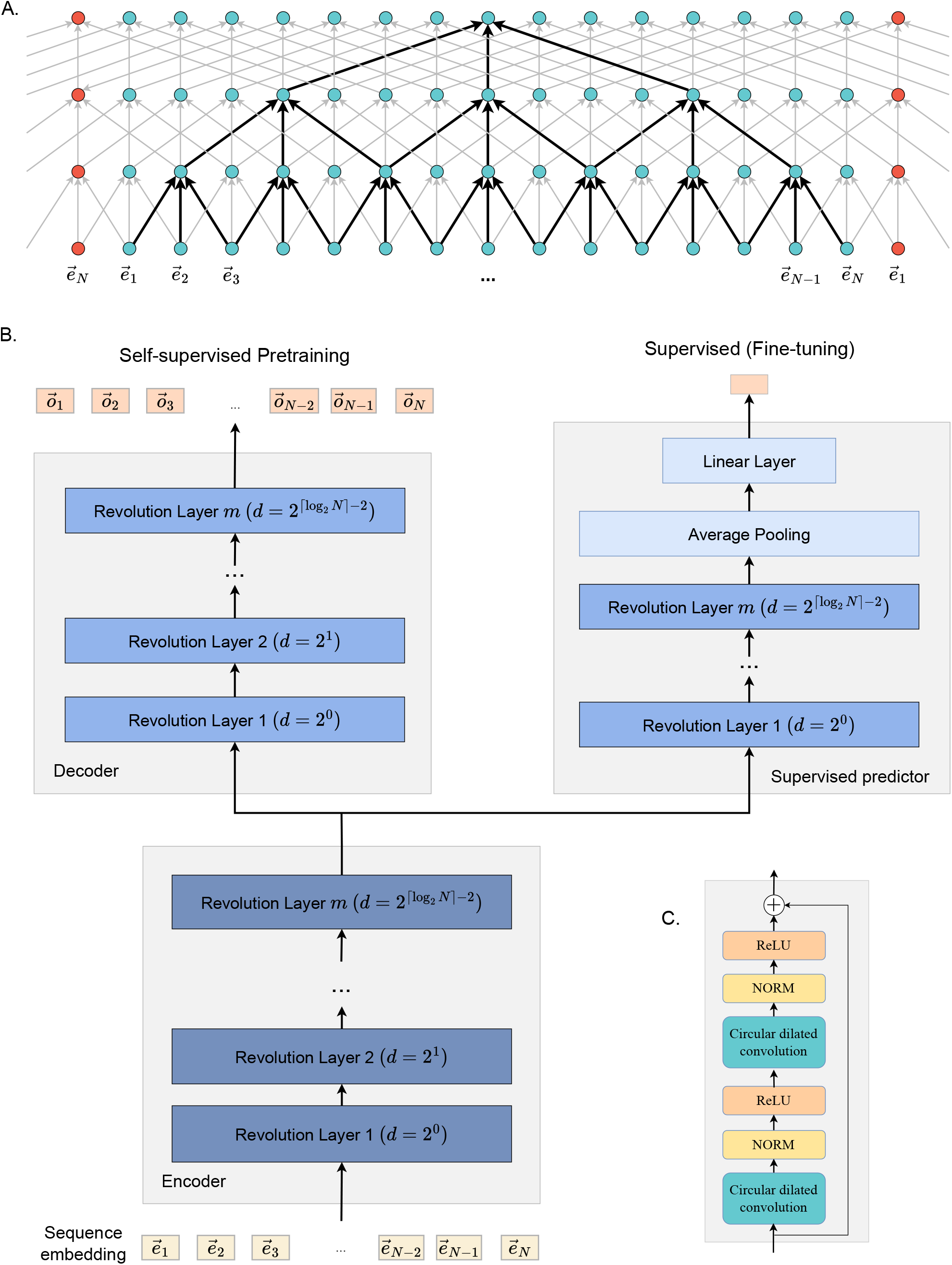
Schematic illustration of Revolution. Panel A shows a Revolution neural network with three layers (dilation={1, 2, 4}, kernel size *K* is 3). The red nodes show how circular padding is applied. Panel B illustrates the self-supervised learning architecture with circular dilated convolution. The self-supervised pretraining branch includes one encoder and one decoder, both being a Revolutional network. The pretrained encoder is then combined with a supervised predictor and fine-tuned with supervised information. After fine-tuning, the right branch (encoder + supervised predictor) is used in sequence-based inference. Panel C presents a single module within one Revolution layer.

A Revolution module contains two circular dilated convolutions, each being followed by batch normalization (NORM) and a nonlinear activation (ReLU). A residual skip connection is used around the operations. The Revolution module is illustrated in Figure 1C.

A Revolution network consists of [log_2_ *N*] − 1 Revolution modules, where the *l*-th module uses dilation *d*_*l*_. Note that we do not shorten the sequence length after each Revolution layer, which is different from typical bottleneck designs in conventional neural networks that contain pooling or low-rank projections. Therefore, Revolution is a wide neural network that directs the input signal toward the learning targets in a broad way.

### Pretraining

Revolution has been applied in our previous work for supervised classification of long sequences (Cheng et al., 2023). This paper shows that Revolution also performs well for self-supervised learning, where all key components in our framework include Revolution networks (see Figure 1B).

Our pretraining follows masked language modeling (MLM; Taylor, 1953), a successful pretraining method in NLP, except that we replace Transformer-like attention modules with Revolution. Our method first maps the original information to a latent representation with an encoder and then reconstructs the masked tokens using a decoder. Because Revolution is good at modeling long-range interactions, we apply it to the constructions of both the encoder and decoder.

In masked learning, we randomly sample a subset of sequence positions as a mask and use the non-masked sequence elements to infer those masked. Similar to BERT, we replace 80% of the selected elements by [0, 0, 0, 0], 10% with the one-hot encoding of a random nucleotide base, ‘A,’ ‘T,’ ‘G,’ or ‘C’, and the rest 10% unchanged.

The input DNA sequence is represented in one-hot encoding, i.e., with four channels. An embedding layer before the encoder maps the sequence elements into a higher dimensional space. Afterward, the sequence embedding 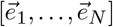 is fed to the encoder (a Revolution network) to obtain its representation in a latent space.

The sequence in latent space is fed to another Revolution network as a decoder which tries to reconstruct the masked bases. Because the reconstruction part appears only in pretraining and is not used in the sequence-based inference, we follow He et al. (2022) and use a lightweight decoder with fewer channels in Revolution. The decoder output 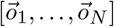 is compared with the original sequence elements at the masked positions.

### Fine-Tuning

After pretraining, the encoder can be used as a feature extractor in downstream sequence-based supervised inference tasks. The process is shown in the right branch of Figure 1B.

The embedding and encoding steps are the same as in pretraining, except that the sequence is not masked. Afterward, the sequence in latent space is fed to a predictor which comprises another Revolution network followed by a position-wise average pooling and a linear layer. The Revolution network in the predictor mixes the encoded information toward the inference target, and the pooling and linear layer perform the final ensemble, e.g., regression or classification. The predictor is trained by supervised learning and possible fine-tuning of the encoder (see Section Ablation Study for details).

### Datasets

We have used genomic data from humans and plants to test Revolution for self-supervised learning and other compared methods. Only DNA sequences are used as input in all experiments.

#### Human sequence data

In the pretraining stage, we extracted a total of 40,000 DNA sequences of 10kbp length from GRCh38 as follows: 1) we randomly selected one of the 24 chromosomes; 2) we randomly selected a starting position on the chosen chromosome; 3) we extracted the DNA sequence of 10kbp from left to right. Each extracted DNA sequence was then one-hot encoded, with the four DNA nucleotide bases A, T, G, C represented as [1, 0, 0, 0], [0, 1, 0, 0], [0, 0, 1, 0], and [0, 0, 0, 1], respectively. We used 80% of the sequences for training, 10% for validation, and 10% for testing.

The downstream binary classification task was to recognize causal genetic variants, i.e., to classify a genetic variant to be likely causal (causal probability > 0.9, as determined by the population-based fine-mapping model SuSiE; Weissbrod et al., 2020) or likely spurious eQTLs (causal probability < 0.01). We have used a binary classification data set prepared by Avsec et al. (2021). Remarkably, the task in our work is more challenging than the original one because it requires direct prediction from DNA sequence to binary class labels, and the gene expression values are unavailable. There are in total 97,992 instances across 49 human tissues.

Each instance includes one reference sequence (from GRCh38), one alternative sequence, and the variant effect label. We randomly split the instances into training (80%), validation (10%), and testing (10%) sets. More details can be found in the supplemental document (Table 1).

**Table 1.**
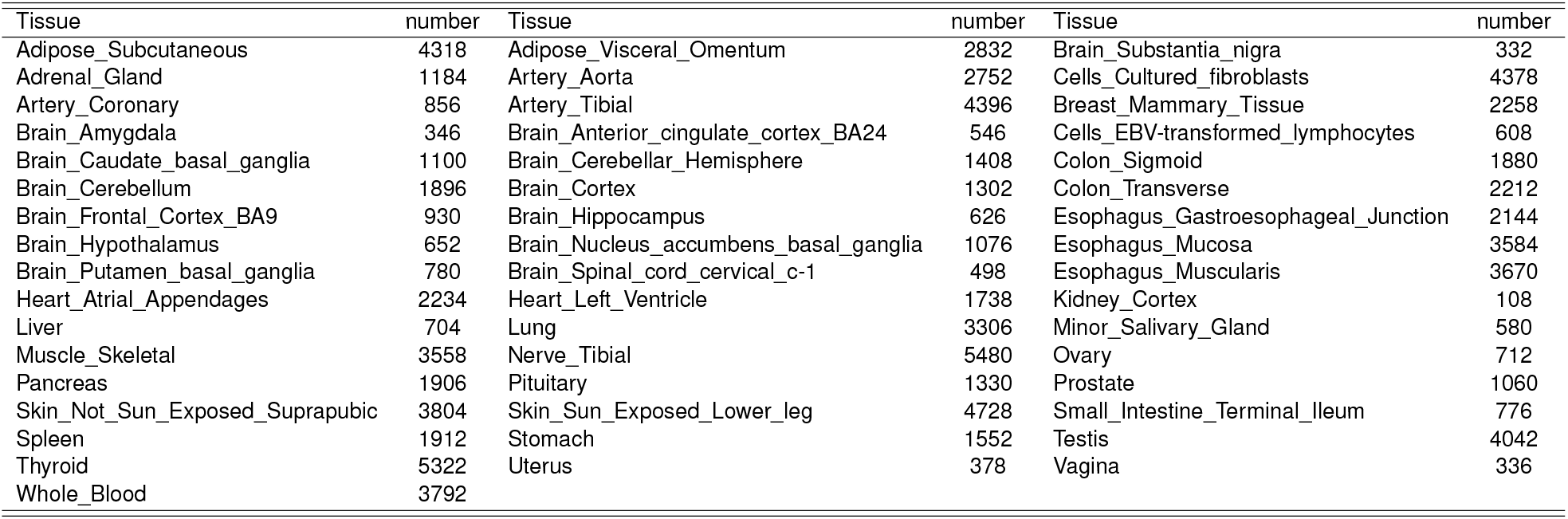
Data number for each GTEx tissue.

**Table 2.**
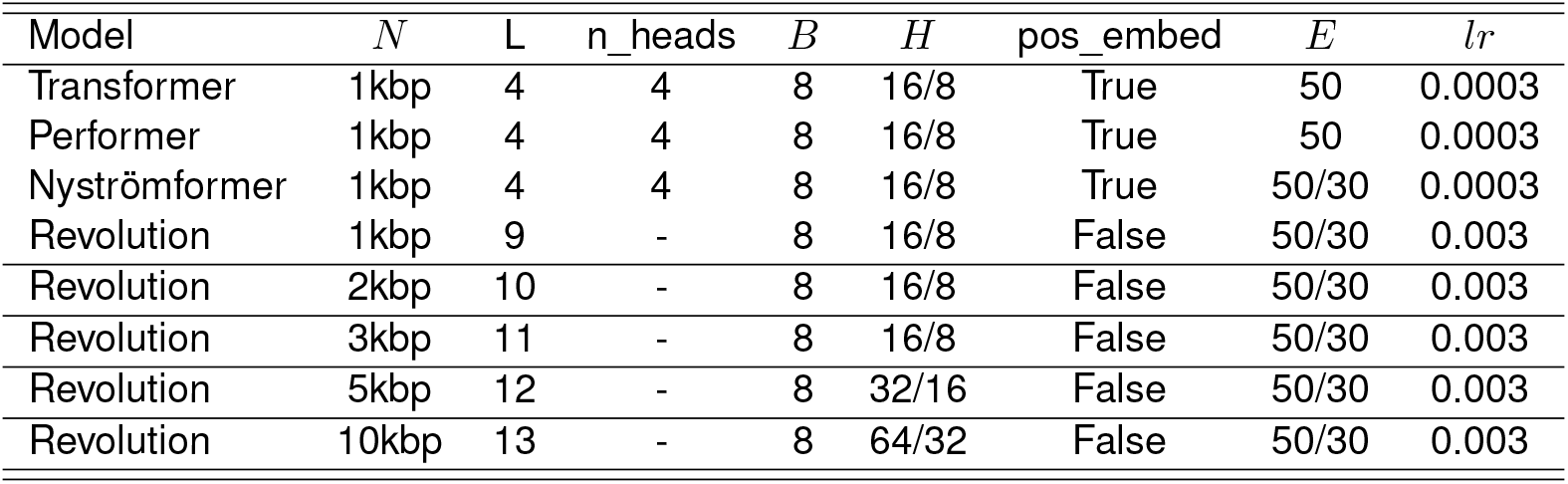
Hyperparameters details for every model on variant effect prediction. *N*, *L*, *B*, *H*, *E*, and *lr* refer to max sequence length, layer number, batch size, hidden states size, running epochs, and learning rate, respectively. The first value of *H* is the dimension of the encoder while the second value is the dimension of the decoder. The two numbers for *E* refer to running epochs of w. pretraining and w/o pretraining.

#### Plant sequence data

The plant genome data set from Zhao et al. (2021) includes six representative species: Arabidopsis thaliana, Brachypodium distachyon, Oryza sativa (rice), Setaria italica, Sorghum bicolor, and Zea mays (maize). We used seven reference genomes, including two rice varieties (O.sativa-MH and O.sativa-ZS) and one genome for each other species.

In the pretraining stage, we extracted 64,000 DNA sequences of 1kbp length from the maize genome and 51,200 sequences from the genomes of the other species for self-supervised training. We also use onehot encodings for the DNA sequences. The data set contains other base symbols, N, M, K, etc., in addition to A, T, G, and C. We encoded these bases as [0.25, 0.25, 0.25, 0.25] following the strategy by Zhao et al. (2021). We pretrained a separate model for each genome variety and obtained seven pretrained models in total.

The downstream binary classification task was to determine whether a given DNA sequence belongs to an open chromatin region (OCR) for a specific feature. OCRs were identified by Assay of Transposase Accessible Chromatin using sequencing (ATAC-seq; Buenrostro et al., 2013). A DNA sequence was labeled as positive if its center 200bp has more than 50% overlap with the OCR, and as negative otherwise. The data set contains multiple features for each plant (ranging from 9 to 19). To evaluate the compared models in few-shot learning, we used only 10% of the supervised labels in training while keeping validation and testing sets unchanged. See the supplemental document (Table 3) for details.

**Table 3.**
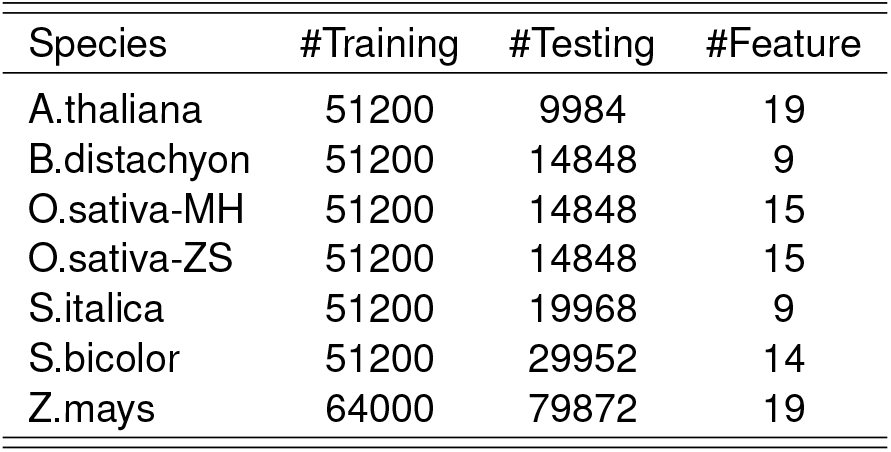
Summary of plant datasets. #Training, #Testing, and #Feature refer to the number of training sequences, validation/testing sequences, and features, respectively.

## Results

### Variant effect prediction in human tissues

Revolution with self-supervised pretraining outperforms the Transformer-like methods in the genetic variant prediction task. Additional hyperparameters are summarized in Supplementary Table 2. Figure 4 left panel shows the comparison between Revolution with popular Transformer variants, Performer and Nyströmformer, for sequence length 1kbp. We can see that Revolution with self-supervised pretraining performs the best with a mean AUROC of 88.08%. Our method is significantly better than the runner-up, Nyströmformer, with *p*-value 0.00021.

Figure 2 presents the comparison with Enformer, the method from the supervised label provider. We can see that all dots are above the dashed line (equal performance), which shows that Revolution with selfsupervised pretraining in general outperforms Enformer. For some tissues, e.g., Brain_Putamen_basal_ganglia and Uterus, the AUROC values are improved by 30.55% and 30.01%, respectively. On average, Revolution increases the AUROC from 70.50% to 91.12%. Because Enformer uses a pyramid design to reduce the sequence length, our win also indicates that it is beneficial to use the original sequence without condensation. Furthermore, we find that pretraining on Revolution helps as the sequence length scales up. We present the AUROC of Revolution (average over all 49 tissues) in Figure 3 for five different sequence lengths. The performance of Revolution improves as the DNA sequence length increases, both with and without pretraining. This result suggests that Revolution can effectively incorporate long-range interactions within DNA sequences. We also observe that Revolution with pretraining performs better than without, indicating that pretraining can benefit Revolution for variant effect prediction. In particular, when the sequence length is 10kbp, pretraining improves the performance by 1.09%, indicating that Revolution can benefit from pretrained information on longer sequences. The best result achieved by Revolution with self-supervised learning is 91.12±0.06% using sequence length 10kbp, which is substantially better than all other existing methods.

**Figure 2.**
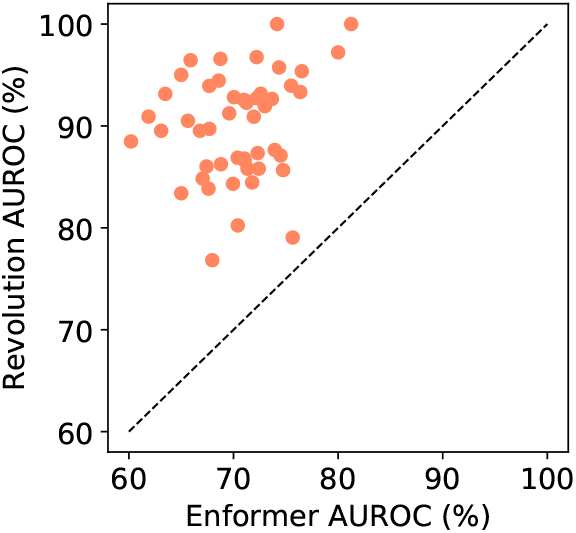
The AUROC *×*100% of Revolution and Enformer on variant effect prediction. Each dot represents a tissue. The dashed line refers to equal performance. A dot above the dashed line means Revolution is better than Enformer for the corresponding tissue. Overall, Revolution is significantly better than Enformer, with a *p*-value < 0.00001 of the *t*-test.

**Figure 3.**
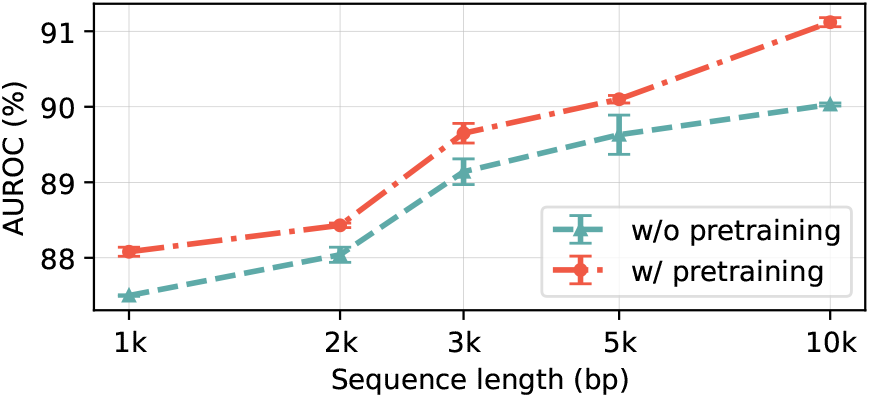
The AUROC *×*100% of Revolution on variant effect prediction with various sequence lengths. Means and standard deviations are computed across five runs.

**Figure 4.**
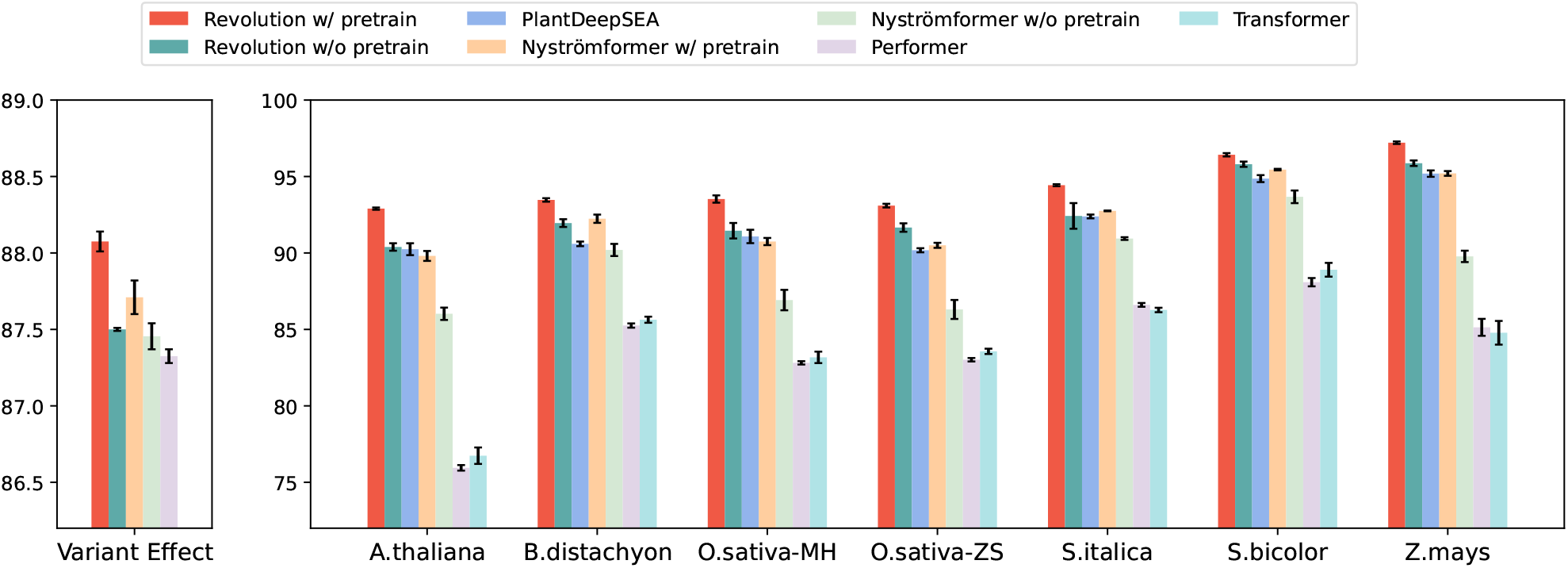
The AUROC ×100% of different models on variant effect (left) and OCR (right) prediction of length 1kbp. Means and standard deviations are computed across five runs. Revolution with pretraining is significantly better than the runner-up for each task, with *p*-values < 0.0005 of the t-test.

### OCR prediction in plants

First, we compared Revolution with other models on sequences of length 1kp. DeepSEA (Zhou and Troyanskaya, 2015) and its deeper version (Chen et al., 2019), which doubles the number of convolutional layers of DeepSEA, are pyramid convolutional architectures designed for human genomic sequences of 1kp. PlantDeepSEA adapts the deeper DeepSEA for plant genomic data. We also compared with Transformer, Performer, and Nyströmformer. More details about the compared methods are given in the supplemental document (Table 4). The performance is evaluated using average AUROC over all features of the plant.

**Table 4.**
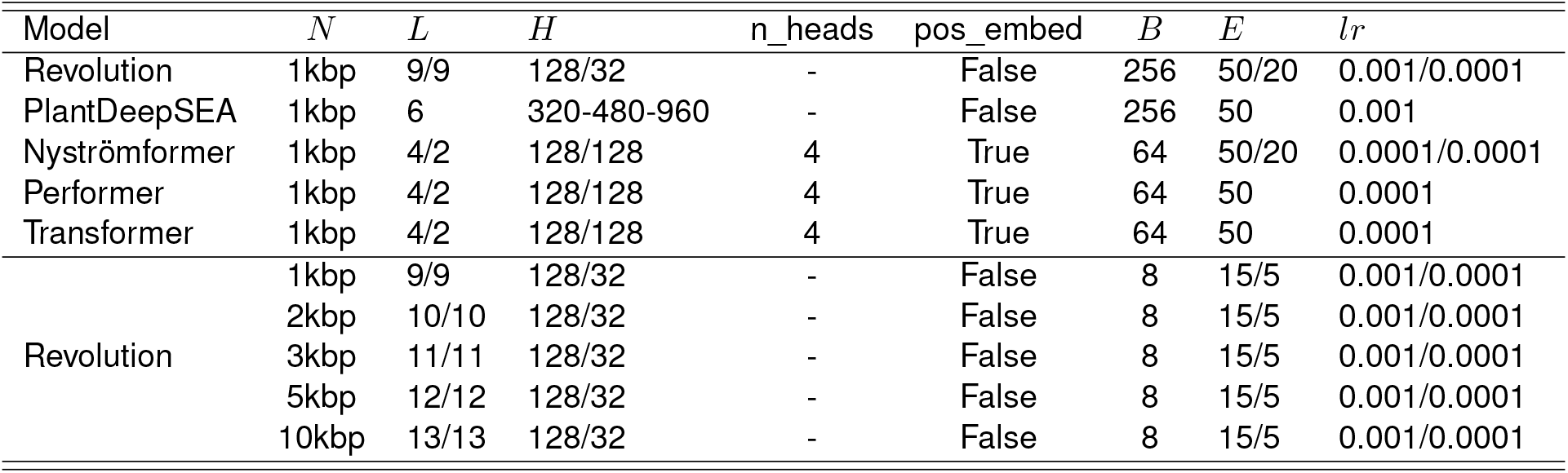
Hyperparameters of every model on OCRs prediction. *N*, *L*, *H*, *B*, *E*, and *lr* refer to the sequence length, the number of layers, hidden states size, batch size, running epochs, and learning rate, respectively. PlantDeepSEA increases the hidden size as layers are stacked. The two parts of *L* and *H* refer to hyperparameters of the Encoder and Decoder or Supervised predictor, respectively. The two parts of *E* and *lr* refer to hyperparameters for without- and with-pretraining, respectively.

Revolution with self-supervised pretraining achieves the best for all plants, as we can see in Figure 4 right panel. The p-values show that our method is significantly better than the runner-ups. Revolution without pretraining is the first runner-up in five of seven cases and comparable to the first runner-up in the remaining.

Next, we studied the performance of Revolution with five different input sequence lengths. The results are shown in Figure 5. We observe similar patterns as in the human tissues: 1) the performance is improving with increasing lengths in most cases, suggesting a wider context often helps OCR predictions in the center 200bp; 2) Revolution benefits from self-supervised pretraining in all cases.

**Figure 5.**
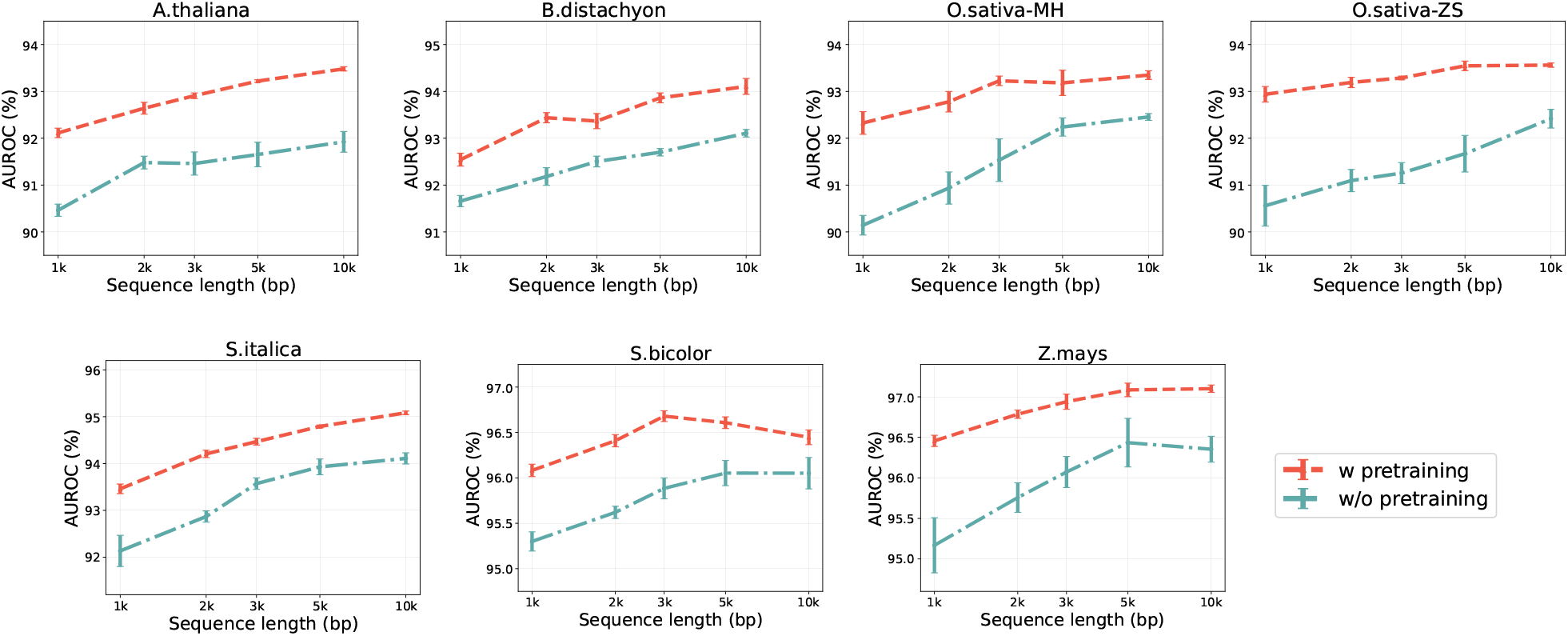
The AUROC *×*100% of Revolution on OCR prediction with increasing sequence lengths. We compared w/o pretraining and w/ pretraining for each plant. Means and standard deviations are computed across five runs.

### Ablation Study

#### Fine-tuning vs. Revolution-probing

In the above experiments, we have used fine-tuning as the default approach for the prediction tasks. Recently, Kumar et al. (2022) suggested that combining linear probing and fine-tuning could yield better results for distribution shift data. We also investigated whether the combination is helpful for DNA sequence-based inferences. Because we used an additional Revolution network on top of the pretrained network during pretraining, we refer to our probing method as Revolution probing (RP) instead of linear probing. We selected a sequence length of 1kbp and compared the performance of three different strategies: fine-tuning (FT), Revolution probing (RP), and a combination of the two (RP-FT). In RP, we froze the pretrained encoder and used it to probe the supervised predictor. In RP-FT, we unfroze the encoder and retrained the whole network, with the supervised predictor initialized with the one obtained in RP.

The investigation result is shown in Figure 6. We can see that RP and RP-FT are better than FT in all cases. Especially, they bring nearly 7% improvement for human genetic variant predictions. RP-FT performs slightly better than RP in some cases. We choose FT instead of RP-FT because the latter requires an extra round of training (RP) and the gain is very small.

**Figure 6.**
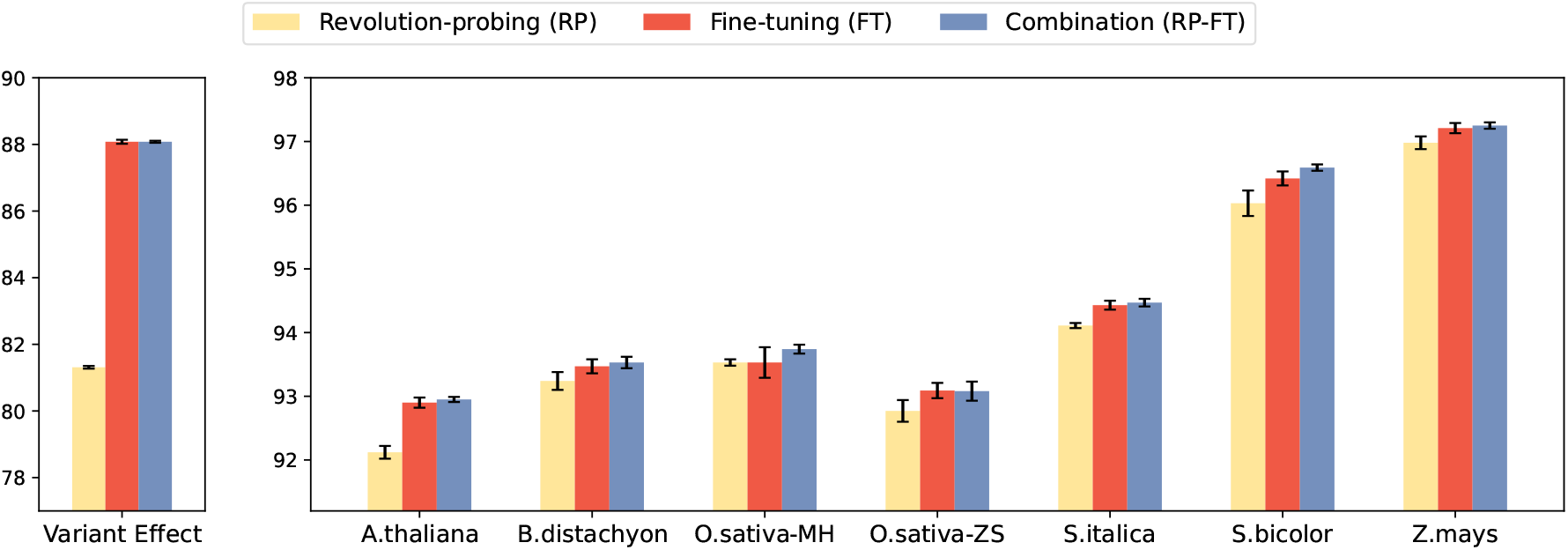
The AUROC *×*100% of Revolution with different supervised learning strategies after pretraining. Means and standard deviations are computed across five runs.

#### Masking Ratio

Masked language models typically use a masking rate of 15%. However, some researchers have also argued that a masking rate of 40% is optimal for large pretraining models (Wettig et al., 2022). Therefore, we investigated how the masking ratio of Revolution influences the performance of DNA sequencebased inferences. We studied a range of different masking ratios from 10% to 90% for both variant effect and OCR prediction tasks. We used sequences of length 1kbp and fixed all the other parameters. The results, as shown in Figure 7, suggest that a lower masking ratio between 10% and 40% is often a good choice, and the optimal ratio can depend on the task and organisms. We empirically selected 30% and 15% for humans and plants, respectively.

**Figure 7.**
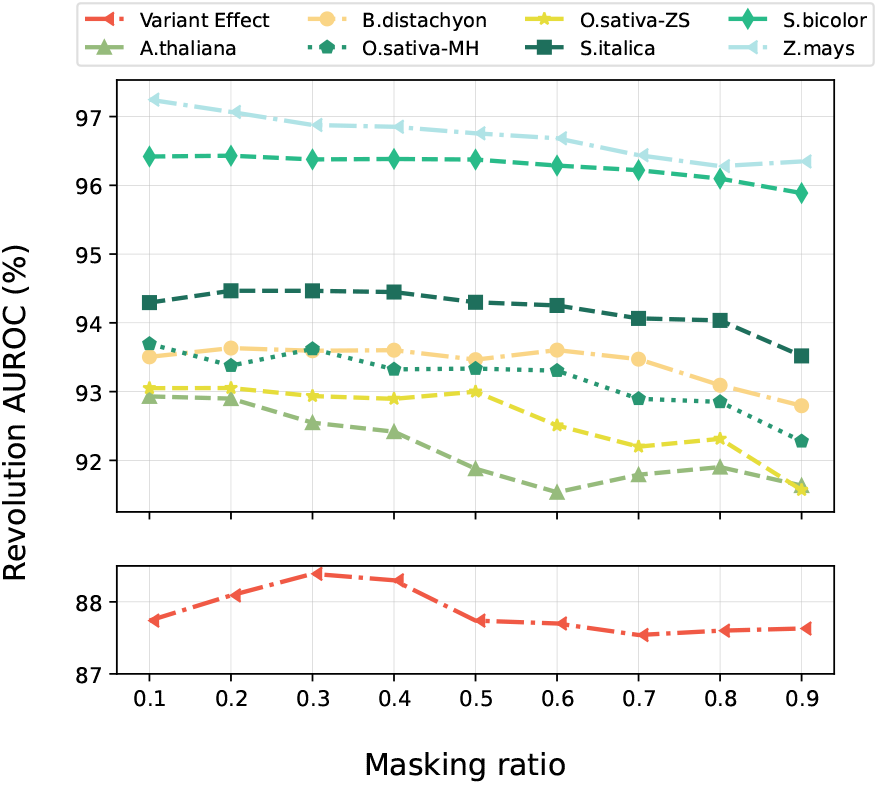
The AUROC *×*100% of Revolution on variant effect and OCR prediction with different masking ratios.

## Discussion

In this paper, we proposed a novel framework called Revolution for sequence-based inferences. Our method has overcome two major challenges in this domain: the inability to handle long DNA sequences and the limited availability of supervised labels. To tackle the first challenge, Revolution uses neural networks as the backbone, which can capture prolonged interactions taking place within DNA sequences. To address the second challenge, Revolution incorporates self-supervised pretraining to improve the model’s capacity. These two design choices enable Revolution to scale up to long DNA sequences and achieve enhanced performance with a few supervised labels. Our experimental results showed that Revolution achieved state-of-the-art performance both on variant effect and OCR prediction tasks. We also observed that the pretrained Revolution model generalized well even when only 1% of the PlantDeepSEA DNA sequences were used. The results also suggested that using fine-tuning and probing together improves the model accuracy. We also studied and suggested an optimal masking ratio for DNA sequence-based MLM for human and plant tissues. Our pretrained Revolution model is freely available on GitHub and can be applied to other sequence-based inference tasks, such as enhancer prediction and taxonomy classification.

While Revolution demonstrates strong learning capabilities, there are still areas for improvement in future work. Some research suggests that pretraining on DNA sequences from different species can benefit downstream tasks more than using DNA from a single species (ref). Therefore, our self-supervised pretraining could be extended to cross-species pretraining to improve the capacity of the pretrained model further. The framework could also be readily applied to other tasks such as predicting gene expressions.

## Acknowledgements

We acknowledge using the IDUN computing cluster (Själander et al., 2019).

## Supplementary Note 1: Human

### A. Data description

In our experiment on variant effect prediction, we use a dataset consisting of 97,992 instances across 49 tissues. The details of the dataset are shown in Table 1. We shuffle the instances and split the data into 80% training, 10% validation, and 10% test sets. The sequences are extracted from Gh38 to construct both the pretraining dataset and the downstream task.

To run Enformer on this dataset, we split each tissue into 80% training, 10% validation, and 10% test sets and use a random forest classifier. This allows us to evaluate the performance of Enformer on the variant effect prediction task.

### B. Hyperparameters

In this work, we focus on predicting variant effects using neural models. Our experiments utilize several hyperparameters, which are shown in Table 1. We maintain a constant batch size across all models and use a mask ratio of 30% during the pretraining stage. Our masking strategy includes random masking, replace masking, and no masking in an 80-10-10 ratio. We run the pretraining for 50 epochs with a learning rate of 0.003.

We use different hidden dimensions for the encoder and decoder according to the sequence length. For example, we use 32 for the encoder and 16 for the decoder when pretraining on 1kbp sequences. The dimension of the decoder is set to half of the encoder to keep it lightweight. In the fine-tuning stage, we run the model for 30 epochs with a learning rate of 0.003. The model first loads the pre-trained encoder and then fine-tunes a Revolution classifier. The hidden dimension of the Revolution classifier is set to half of the pre-trained encoder.

## Supplementary Note 2: Plants

### A. Data description

In our experiment on OCRs prediction, we only use 1% of labeled training sequences to show the benefit of self-supervised learning when supervised labels are limited. Following the strategy of Zhao et al. (2021), we select 1 or 2 chromosomes as the validation or test set and exclude them from the train set. Each plant predicts OCRs in different tissues and thus has multiple tasks. Table 3 summarizes the datasets of different plants.

### B. Hyperparameters

All plants achieved the best average AUROC on downstream tasks when the masking ratio is either 10% or 20%, and we thus selected 15% for all plants for simplicity. Random masking, replace masking, and no masking followed an 80-10-10 split. We ran pretraining for 50 epochs with the learning rate of 0.001 and 0.0001 for Revolution and Nyströmformer, respectively. More hyperparameters are shown in Table 4.

