## Supplemental Tables for "Self-supervised learning for DNA sequences with circular dilated convolutional networks"

### Supplementary Note 1: Human

#### A. Data description

In our experiment on variant effect prediction, we use a dataset consisting of 97,992 instances across 49 tissues. The details of the dataset are shown in Table 1. We shuffle the instances and split the data into 80% training, 10% validation, and 10% test sets. The sequences are extracted from Gh38 to construct both the pretraining dataset and the downstream task.

To run Enformer on this dataset, we split each tissue into 80% training, 10% validation, and 10% test sets and use a random forest classifier. This allows us to evaluate the performance of Enformer on the variant effect prediction task.

**Table 1.** Data number for each GTEx tissue.

| Tissue | number | Tissue | number | Tissue | number |
|---|---|---|---|---|---|
| Adipose_Subcutaneous | 4318 | Adipose_Visceral_Omentum | 2832 | Brain_Substantia_nigra | 332 |
| Adrenal_Gland | 1184 | Artery_Aorta | 2752 | Cells_Cultured_fibroblasts | 4378 |
| Artery_Coronary | 856 | Artery_Tibial | 4396 | Breast_Mammary_Tissue | 2258 |
| Brain_Amygdala | 346 | Brain_Anterior_cingulate_cortex_BA24 | 546 | Cells_EBV-transformed_lymphocytes | 608 |
| Brain_Caudate_basal_ganglia | 1100 | Brain_Cerebellar_Hemisphere | 1408 | Colon_Sigmoid | 1880 |
| Brain_Cerebellum | 1896 | Brain_Cortex | 1302 | Colon_Transverse | 2212 |
| Brain_Frontal_Cortex_BA9 | 930 | Brain_Hippocampus | 626 | Esophagus_Gastroesophageal_Junction | 2144 |
| Brain_Hypothalamus | 652 | Brain_Nucleus_accumbens_basal_ganglia | 1076 | Esophagus_Mucosa | 3584 |
| Brain_Putamen_basal_ganglia | 780 | Brain_Spinal_cord_cervical_c-1 | 498 | Esophagus_Muscularis | 3670 |
| Heart_Atrial_Appendages | 2234 | Heart_Left_Ventricle | 1738 | Kidney_Cortex | 108 |
| Liver | 704 | Lung | 3306 | Minor_Salivary_Gland | 580 |
| Muscle_Skeletal | 3558 | Nerve_Tibial | 5480 | Ovary | 712 |
| Pancreas | 1906 | Pituitary | 1330 | Prostate | 1060 |
| Skin_Not_Sun_Exposed_Suprapubic | 3804 | Skin_Sun_Exposed_Lower_leg | 4728 | Small_Intestine_Terminal_Ileum | 776 |
| Spleen | 1912 | Stomach | 1552 | Testis | 4042 |
| Thyroid | 5322 | Uterus | 378 | Vagina | 336 |
| Whole_Blood | 3792 | | | | |

#### B. Hyperparameters

In this work, we focus on predicting variant effects using neural models. Our experiments utilize several hyperparameters, which are shown in Table 1. We maintain a constant batch size across all models and use a mask ratio of 30% during the pretraining stage. Our masking strategy includes random masking, replace masking, and no masking in an 80-10-10 ratio. We run the pretraining for 50 epochs with a learning rate of 0.003.

We use different hidden dimensions for the encoder and decoder according to the sequence length. For example, we use 32 for the encoder and 16 for the decoder when pretraining on 1kbp sequences. The dimension of the decoder is set to half of the encoder to keep it lightweight. In the fine-tuning stage, we run the model for 30 epochs with a learning rate of 0.003. The model first loads the pre-trained encoder and then fine-tunes a Revolution classifier. The hidden dimension of the Revolution classifier is set to half of the pre-trained encoder.

**Table 2.** Hyperparameters details for every model on variant effect prediction. $N$, $L$, $B$, $H$, $E$, and $lr$ refer to max sequence length, layer number, batch size, hidden states size, running epochs, and learning rate, respectively. The first value of $H$ is the dimension of the encoder while the second value is the dimension of the decoder. The two numbers for $E$ refer to running epochs of w. pretraining and w/o pretraining.

| Model | $N$ | L | n_heads | $B$ | $H$ | pos_embed | $E$ | $lr$ |
|---|---|---|---|---|---|---|---|---|
| Transformer | 1kbp | 4 | 4 | 8 | 16/8 | True | 50 | 0.0003 |
| Performer | 1kbp | 4 | 4 | 8 | 16/8 | True | 50 | 0.0003 |
| Nyströmformer | 1kbp | 4 | 4 | 8 | 16/8 | True | 50/30 | 0.0003 |
| Revolution | 1kbp | 9 | - | 8 | 16/8 | False | 50/30 | 0.003 |
| Revolution | 2kbp | 10 | - | 8 | 16/8 | False | 50/30 | 0.003 |
| Revolution | 3kbp | 11 | - | 8 | 16/8 | False | 50/30 | 0.003 |
| Revolution | 5kbp | 12 | - | 8 | 32/16 | False | 50/30 | 0.003 |
| Revolution | 10kbp | 13 | - | 8 | 64/32 | False | 50/30 | 0.003 |

### Supplementary Note 2: Plants

#### A. Data description

In our experiment on OCRs prediction, we only use 1% of labeled training sequences to show the benefit of self-supervised learning when supervised labels are limited. Following the strategy of Zhao et al. (2021), we select 1 or 2 chromosomes as the validation or test set and exclude them from the train set. Each plant predicts OCRs in different tissues and thus has multiple tasks. Table 3 summarizes the datasets of different plants.

**Table 3.** Summary of plant datasets. #Training, #Testing, and #Feature refer to the number of training sequences, validation/testing sequences, and features, respectively.

| Species | #Training | #Testing | #Feature |
|---|---|---|---|
| A.thaliana | 51200 | 9984 | 19 |
| B.distachyon | 51200 | 14848 | 9 |
| O.sativa-MH | 51200 | 14848 | 15 |
| O.sativa-ZS | 51200 | 14848 | 15 |
| S.italica | 51200 | 19968 | 9 |
| S.bicolor | 51200 | 29952 | 14 |
| Z.mays | 64000 | 79872 | 19 |

##### B. Hyperparameters

All plants achieved the best average AUROC on downstream tasks when the masking ratio is either 10% or 20%, and we thus selected 15% for all plants for simplicity. Random masking, replace masking, and no masking followed an 80-10-10 split. We ran pretraining for 50 epochs with the learning rate of 0.001 and 0.0001 for Revolution and Nyströmformer, respectively. More hyperparameters are shown in Table 4.

**Table 4.** Hyperparameters of every model on OCRs prediction. $N$, $L$, $H$, $B$, $E$, and $lr$ refer to the sequence length, the number of layers, hidden states size, batch size, running epochs, and learning rate, respectively. PlantDeepSEA increases the hidden size as layers are stacked. The two parts of $L$ and $H$ refer to hyperparameters of the Encoder and Decoder or Supervised predictor, respectively. The two parts of $E$ and $lr$ refer to hyperparameters for without- and with- pretraining, respectively.

| Model | $N$ | $L$ | $H$ | n_heads | pos_embed | $B$ | $E$ | $lr$ |
|---|---|---|---|---|---|---|---|---|
| Revolution | 1kbp | 9/9 | 128/32 | - | False | 256 | 50/20 | 0.001/0.0001 |
| PlantDeepSEA | 1kbp | 6 | 320-480-960 | - | False | 256 | 50 | 0.001 |
| Nyströmformer | 1kbp | 4/2 | 128/128 | 4 | True | 64 | 50/20 | 0.0001/0.0001 |
| Performer | 1kbp | 4/2 | 128/128 | 4 | True | 64 | 50 | 0.0001 |
| Transformer | 1kbp | 4/2 | 128/128 | 4 | True | 64 | 50 | 0.0001 |
| Revolution | 1kbp | 9/9 | 128/32 | - | False | 8 | 15/5 | 0.001/0.0001 |
| | 2kbp | 10/10 | 128/32 | - | False | 8 | 15/5 | 0.001/0.0001 |
| | 3kbp | 11/11 | 128/32 | - | False | 8 | 15/5 | 0.001/0.0001 |
| | 5kbp | 12/12 | 128/32 | - | False | 8 | 15/5 | 0.001/0.0001 |
| | 10kbp | 13/13 | 128/32 | - | False | 8 | 15/5 | 0.001/0.0001 |
